# The *Vibrio cholerae* Seventh Pandemic islands act in tandem to defend against a circulating phage

**DOI:** 10.1101/2022.05.16.492052

**Authors:** Brendan J O’Hara, Munirul Alam, Wai-Leung Ng

## Abstract

The current circulating pandemic El Tor biotype of *Vibrio cholerae* has persisted for over sixty years and is characterized by its acquisition of two unique genomic islands called the *Vibrio* Seventh Pandemic Islands 1 and 2 (VSP-I and VSP-II). However, the functions of most of the genes on VSP-I and VSP-II are unknown and the advantages realized by El Tor through these two islands are not clear. Recent studies have broadly implicated these two mobile genetic elements with phage defense. Still, protection against phage infection through these islands has not been observed directly in any *V. cholerae* El Tor biotype. Here we report the isolation of a circulating phage from a cholera patient stool sample and demonstrate that propagation of this phage in its native host is inhibited by elements in both VSP-I and VSP-II, providing direct evidence for the role of these genomic islands in phage defense. Moreover, we show that these defense systems are regulated by quorum sensing and active only at certain cell density. Finally, we have isolated a naturally occurring phage variant that is resistant to the defense conferred by the VSP islands, illustrating the countermeasures used by phages to evade these defense mechanisms. Together, this work demonstrates a functional role for the VSPs in *V. cholerae* and highlights the key regulatory and mechanistic insights that can be gained by studying anti-phage systems in their native contexts.

**SIGNIFICANCE (AUTHOR SUMMARY):** The current pandemic strain of *Vibrio cholerae* carries two unique genomic islands. How these two islands confer evolutionary advantage to the pathogen is unknown. We show here the identification of a circulating phage that is sensitive to the defense systems present on these two islands and demonstrate how phage variants can evade these defenses. Our studies provide the first direct evidence showing the importance of these genomic islands in defending against phage in their native environments; and in doing so provide novel insight into the mechanisms of these highly conserved defense elements.

## INTRODUCTION

There have been seven recorded cholera pandemics caused by two *V. cholerae* biotypes: classical and El Tor. The current, longest lasting, (7^th^) cholera pandemic is caused by the El Tor biotype which has globally displaced the classical biotype in both endemic and clinical populations [1]. One of the defining genetic features of this 7^th^ pandemic strain is the presence of the two aptly named *Vibrio* Seventh Pandemic Islands (VSP-I and VSP-II [2]). In spite of their persistence in *V. cholerae*, the functions of most of the genes on VSP-I and VSP-II are unknown and the evolutionary advantages realized by the El Tor biotype through the acquisition of these two islands are not clear.

It is generally believed that concentrations of phage and *V. cholerae* inversely correlate in aquatic reservoirs, and this predator-prey relationship has been postulated to be one driving force in determining the severity and timing of cholera outbreaks [3, 4]. In the last decade, three predominant lineages of virulent *V. cholerae* phages, called ICP1, ICP2, and ICP3, were isolated from Bangladeshi clinical samples [5]. While there is strong evidence for phage predation in cholera patients, only ICP1 was able to prey on *V. cholerae* in estuarine water. ICP2 and ICP3 are better adapted for predation in a nutrient rich environment [6]. A phage cocktail comprised of these three phages is efficient at killing *V. cholerae* both *in vitro* and *in vivo* [7]. While using phages to treat and prevent cholera is promising, the interaction between these *V. cholerae* phages and the bacterial host is often complex and a constant arms race exists between phages and their bacterial hosts [8]. Bacteria can acquire various phage resistance mechanisms through mutations and horizontal gene transfer. Conversely, due to a strong selective pressure, phages readily develop resistance to many bacterial phage defense systems [9]. Therefore, it is important to study phage-pathogen interactions to have a better understanding of transmission of disease, acquisition of new traits important for pathogenesis, and ultimately the evolutionary history of the cholera pandemics.

Recent studies have increasingly shown a connection between predicted phage defense genes and mobile genetic elements (MGEs) [9, 10]. While there has been strong evidence that dedicated anti-phage islands such as the PLE [11] are present in *V. cholerae*, it is apparent that even previously well described MGEs frequently contain phage defense genes. For example, the *V. cholerae* integrative and conjugative element (ICE) SXT has been shown to carry an arsenal of anti-phage gene clusters [12] in addition to the antibiotic resistance genes that were initially characterized. Similarly, variants of the Vibrio Pathogenicity Island I (VPI-I) have been sequenced and found to contain anti-phage CRISPR systems [13].

Until recently, the only VSP-I genes with a described function are *dncV* and *capV*. Respectively these genes encode the cyclic-GMP-AMP(cGAMP) synthase[14] and the cGAMP sensing phospholipase [15]. When the native *capV-dncV* operon from *V. cholerae* is transplanted into a phage-sensitive *E. coli* strain, the recipient strain becomes resistant to certain phages [16]. Related cyclic-oligonucleotide synthase and effector pairs have been subsequently discovered and some were shown to have phage defense functions [17]. Collectively, these anti-phage systems are known as cyclic oligonucleotide-based antiphage signaling systems (CBASS) [18]. Moreover, it is now apparent that the first gene of the VSP-I island, *vc0175,* encodes a functional deoxycytidine deaminase named AvcD [19, 20]. AvcD converts dCMP/dCTP to dUMP/dUTP and alters the cellular nucleotide pools, which is predicted to negatively impact highly replicative elements including phages. Similar to CBASS, AvcD has been demonstrated to inhibit phage when expressed in *E. coli* [19]. So far, neither DncV/CapV nor AvcD has been shown to defend against any phage that infects *V. cholerae* where the system is natively found.

VSP-II is the larger of the two islands and similar to VSP-I it is only in recent years that its function is being examined. Study of this island is complicated by the fact that multiple circulating variants exist [21]. However, most standard *V. cholerae* laboratory strains appear to contain the complete repertoire of VSP-II genes. For years only the island’s integrase and cognate recombination directionality factor were known [22]. It was recently discovered that a gene cluster in VSP-II (*vc0513-5015,*) is a part of the zinc-dependent Zur regulon [23]. Specifically, *vc0513* encodes a transcriptional activator VerA to activate expression of *vc0512* (*aerB)* to alter chemotaxis and aggregation in an oxygen dependent manner.

During the current study, a gene cluster on VSP-II was identified for defense against different MGEs [24]. It is suggested that the *vc0490-492* operon (*ddmABC*) in VSP-II encodes a defense system to reduce plasmid stability and defend against phage. This novel finding greatly increases our understanding of why seventh pandemic El tor biotypes have maintained VSP islands in their genome and why plasmids have been remarkably unstable in these strains. However, while plasmid destabilization by VC0490-492 (DdmABC) was comprehensively shown in *V. cholerae*, phage defense was demonstrated via ectopic expression in *E. coli* and no *Vibriophage* has yet been identified that is susceptible to such a system [24].

Here we report the isolation of a circulating variant of the phage ICP3 from a cholera patient stool sample and demonstrate that propagation of this phage is inhibited by elements independently identified in both VSP-I and VSP-II, providing the first direct evidence on the roles of these genomic islands in phage defense in their native host. We further demonstrate that the defense element on VSP-II is controlled via quorum sensing, providing protection at specific cell densities. Moreover, by comparing the phages that are either resistant or sensitive to the VSP islands, we gain insight on the potential mechanism for evading these systems.

## RESULTS

### Isolation of VSP-I/VSP-II susceptible phage

Recent studies suggest phage defense systems are often found in mobile genetic elements present in bacterial genomes [25]. We therefore suspected that the *Vibrio* Seventh Pandemic Islands (i.e., VSP-I and VSP-II, **Fig.1A**) may also harbor anti-phage genes. However, so far, no *V. cholerae* phage has been reported to be sensitive to these two islands. Due to the prevalence of these two genomic islands in the circulating clinical isolates of *V. cholerae* [2,26,27], we hypothesize that novel phages that are sensitive to these islands exist and can be isolated. Therefore, we screened for novel phages from rice water stool (RWS) samples that had been collected from cholera patients in Bangladesh on permissive host, which is an O1 El Tor *V. cholerae* E7946 derivative lacking both VSP-I and VSP-II islands (ΔVSP) and deleted for prophages CTXΦ and K139 [28, 29] . Frozen RWS samples that have been stored at -80°C were thawed, bacteria and other debris were gently pelleted, and the supernatant was filter sterilized. The filtrate was then plated on soft agar plates containing the ΔVSP host (a general phage isolation and identification scheme is shown in **Fig. 1B**). Four RWS samples were examined, two of these samples contained no detectable plaques while the other two yielded myriad plaques on this particular *V. cholerae* host. These phage plaques were then picked for further examination. It has previously been shown that there are three dominant phages (ICP1, ICP2, and ICP3) circulating with *V. cholerae* in Bangladesh [5]. Using PCR primers specific to the gene encoding the unique DNA polymerase of each phage, we were able to categorize most of the newly isolated phage as closely related to ICP1, ICP2, or ICP3 **(Supplemental Table 1).**

**Figure 1:**
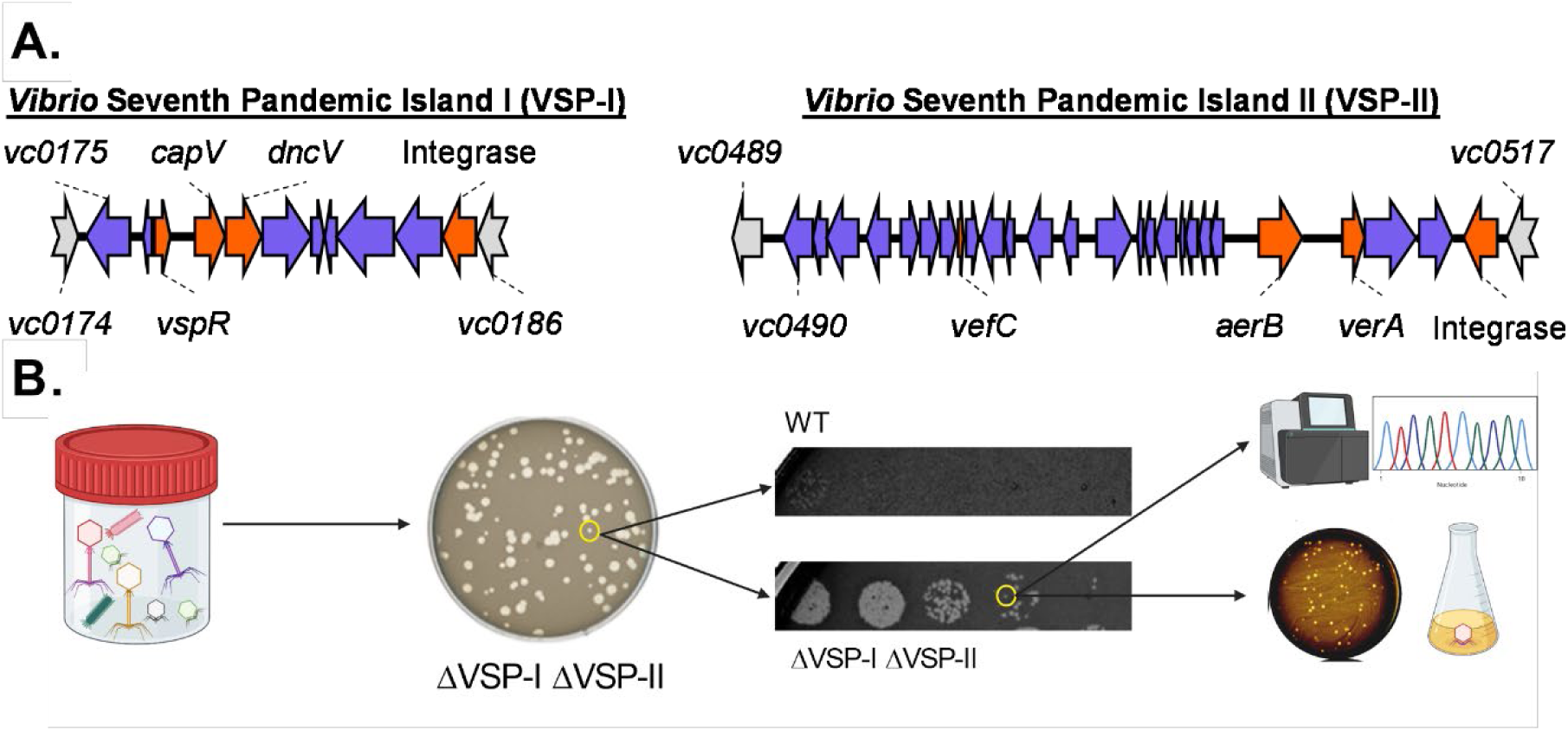
Isolation of VSP-sensitive phages from cholera patient stool samples. (A) Genomic organization of VSP-I &VSP-II, drawn to scale. Genes characterized before this study are marked in orange. Genes on either island with unknown function are blue. Non-VSP genes are grey. (B) Workflow for isolation of VSP sensitive phage. Filter sterilized supernatant from stool samples was plated on a lawn of the ΔVSP-I ΔVSP-II mutant of *V. cholerae.* Plaques were picked, diluted and spot plated on multiple hosts to identify changes in plaquing. Plaques were picked and plaque purified before sequencing and further downstream assays. This figure is created with Biorender.com.

The newly isolated phages were serially diluted and spotted on lawns of either the parental strain (WT) or ΔVSP (**Fig. 1B**). Comparing the plaquing of these phages, we observed the majority showed no host preference. However, a number of phages from one stool sample appeared to form fewer and smaller plaques on WT than on ΔVSP (**Fig. 1B and 2A**). In addition to displaying distinct plaque morphology on each host (**Fig. 2A**), quantitatively, these differentially plaquing phages formed ∼2.5 times fewer plaques on WT than on ΔVSP (**Fig. 2B**). While plaque morphology or efficiency of plating (EOP) can be influenced by many factors, our results suggest that the VSPs play an important role in altering the phage lifecycle. Based on whole genome sequencing, these novel phages all appeared to be variants of ICP3 (Accession number ON464735). We renamed one of these newly isolated VSP susceptible phage ICP3_2016_M1 (M1Φ) since it was isolated from a 2016 stool sample from Bangladesh. ICP3 is a roughly 38kb T7-like lytic phage with a characteristic podoviridae appearance [5]. Compared to previously sequenced ICP3 (ICP3_2009_A), M1Φ has homology with 99% coverage and 95.5% identity. It should be noted that ICP3_2009_A was previously isolated using VSP-carrying El Tor strain [5]. Despite the similarities to previously sequenced ICP3, the significant number of polymorphisms in the M1Φ genome make it difficult to determine any one specific factor responsible for sensitivity to the VSPs that is not observed in ICP3_2009_A (**Supplemental Fig. 1A**).

**Figure 2:**
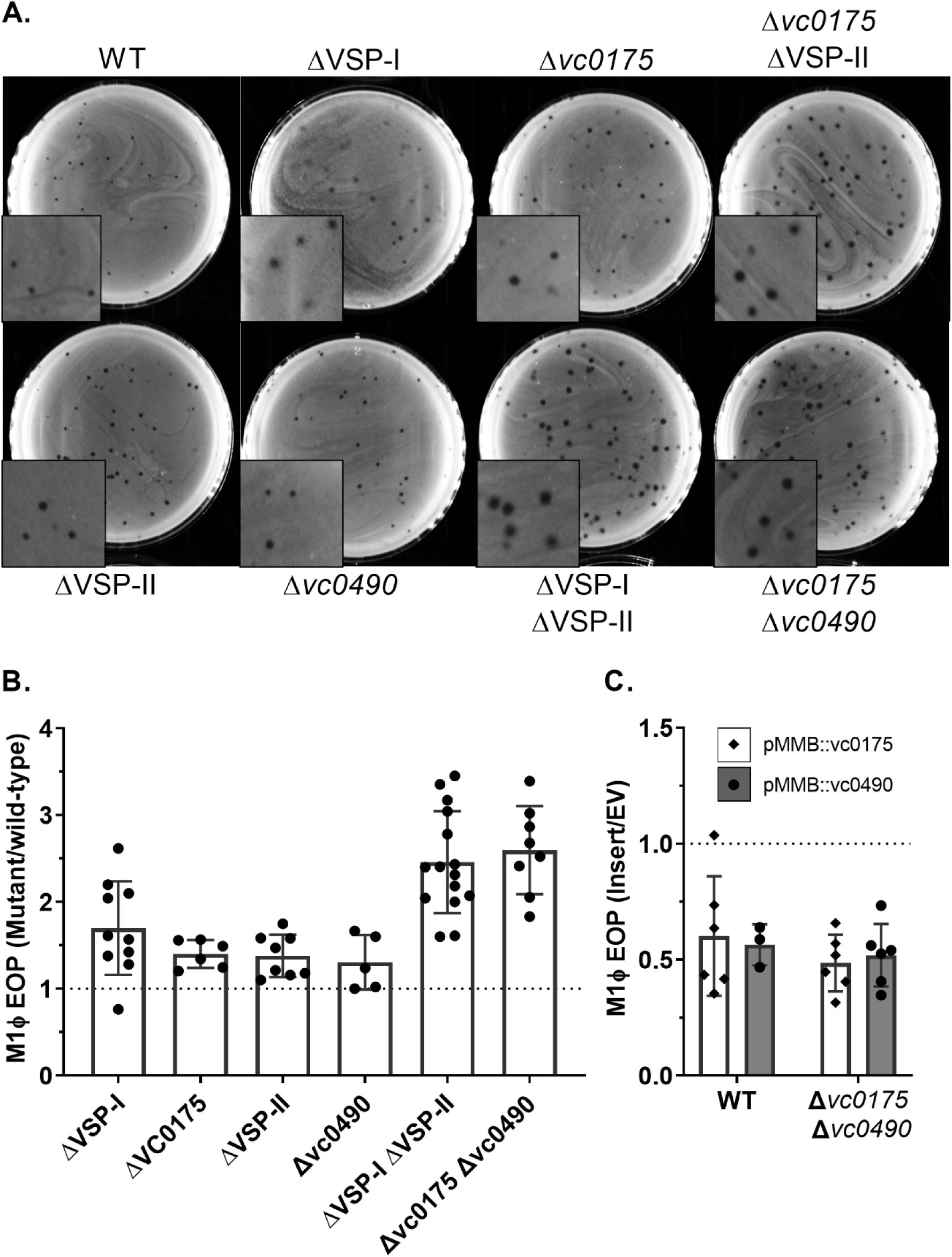
VC0175 (VSP-I) and VC0490 (VSP-II) inhibit phage plaque formation. (A) Representative images of M1Φ plaqued on 0.7% LB agar with various hosts. (B) Efficiency of plating (EOP) of M1Φ on the indicated hosts vs WT. EOP was calculated by dividing the number of plaques obtained from the indicated mutant strain by the plaques obtained from the parental VSP^+^ strain (WT). (C) EOP of different *V. cholerae* strains expressing *vc0175* or *vc0490* from a plasmid. Strains containing plasmid expressing v*c0175 (*white bars/diamonds) or plasmid expressing *vc0490* (grey bars/circles), or plasmid with no insert (EV), on the low copy number plasmid pMMB67eh were infected with M1Φ under inducing conditions in either the WT or Δ*vc0175* Δ*vc0490* background as indicated in the figure. EOP was calculated by dividing the number of plaques from the strains ectopically expressing either *vc0175* or *vc0490* by the number of plaques from the strains carrying the empty vector.

### Determination of VSP genes responsible for M1Φ targeting

In strains with either VSP-I or VSP-II completely deleted, plaque morphology and EOP of M1Φ were modestly changed and more closely resembled what was observed on WT (**Fig. 2A and B**). This implies that there is an element on each island that can act on M1Φ. Looking for homology to other anti-phage systems, we were unable to come up with strong candidates on VSP-II. However, VSP-I contains *dncV*, which encodes the enzyme for 3’-3’-cyclic-GMP-AMP (cGAMP) synthesis[30]. cGAMP activates the phospholipase encoded by the upstream gene *capV* leading to cell death[15]. This *dncV-capV* system had been expressed heterologously in *E. coli* to defend against phage infection [16] and therefore appeared a strong candidate for the VSP-I element for targeting M1Φ. However, we did not observe any differences in either plaque number or morphology when these genes were disrupted with and without VSP-II **(Supplemental Fig. 1B-C).**

To determine the specific VSP gene responsible for altering M1Φ infection, we took a systematic approach to determine the potential anti-phage element(s) on each island. A series of strains were constructed where a defined section of VSP was deleted in a host where the other island had been completely deleted. The regions selected for deletion are loosely based around the predicted operon structure and therefore vary in size. By plaquing M1Φ on these newly constructed “scanning deletion” strains and comparing the number of plaques to that on WT, we identified the regions on each island that appeared to target M1Φ. Specifically, we determined that the plaque number and morphology of M1Φ on the Δ*vc0175-176* ΔVSP-II and ΔVSP-I Δ*vc0490-493* strains were similar to those on the ΔVSP strain (**Supplemental Fig. 1C-D**). To further narrow down the gene on each island, we assayed M1Φ plaque formation on strains with deletions within these newly identified regions. Through this process, we determined that *vc0175* and *vc0490* were the two genes on VSP-I and VSP-II, respectively, which cause differential M1Φ plaquing (**Fig. 2B** and **Supplemental Fig. 1C-E**). Importantly, deletion of just these two genes together phenocopied the deletion of the entirety of both VSP-I (14kb) and VSP-II (27kb). Constitutive expression of either gene from a plasmid in the Δ*vc0175* Δ*vc0490* background reduces plaquing to approximately half of that of an empty vector in the same strain (**Fig. 2** C), which is the exact difference we observed comparing the two single deletion mutants to the double deletion mutants (**Fig. 2B**). Moreover, constitutive expression of either *vc0175* or *vc0490* in WT reduced plaquing as well, suggesting that these two phage defense proteins are not self-limiting and are not subjected to a feedback inhibition mechanism (**Fig. 2C**). From these data we conclude that *vc0175* and *vc0490* are the elements on VSP-I and VSP-II that inhibit the ability of M1Φ to plaque on *V. cholerae.* A recent report indicates *vc0490* forms an operon with *vc0491* and *vc0492* [24], and the three genes function together as a complex, therefore, we assume disruption of any one of the genes in this operon is sufficient to abolish phage defense.

### VC0175 and VC0490 inhibit phage replication

Having established the minimal VSP inhibitory elements, we began to characterize how these genes affect the phage life cycle in its *V. cholerae* host. Deletions of these genes in *V. cholerae* did not inhibit growth (**Fig 3A**) which allowed us to examine the kinetics of M1Φ phage infection across different hosts. To do this we grew cells to mid-log phase and infected a multiplicity of infection (MOI) of 2, indicating there are an average of two phage for every bacterial cell. In contrast to the modest EOP differences observed when plaquing on a plate, the changes in lysis between strains was striking (**Fig. 3A**). The parental VSP^+^ strain (WT) was nearly fully protected from lysis by M1Φ while the Δ*vc0175* Δ*vc0490* mutant was rapidly lysed by the phage. These results directly demonstrate these genes provide a high level protection to a rapidly growing bacterial population against phage predation in liquid culture.

**Figure 3:**
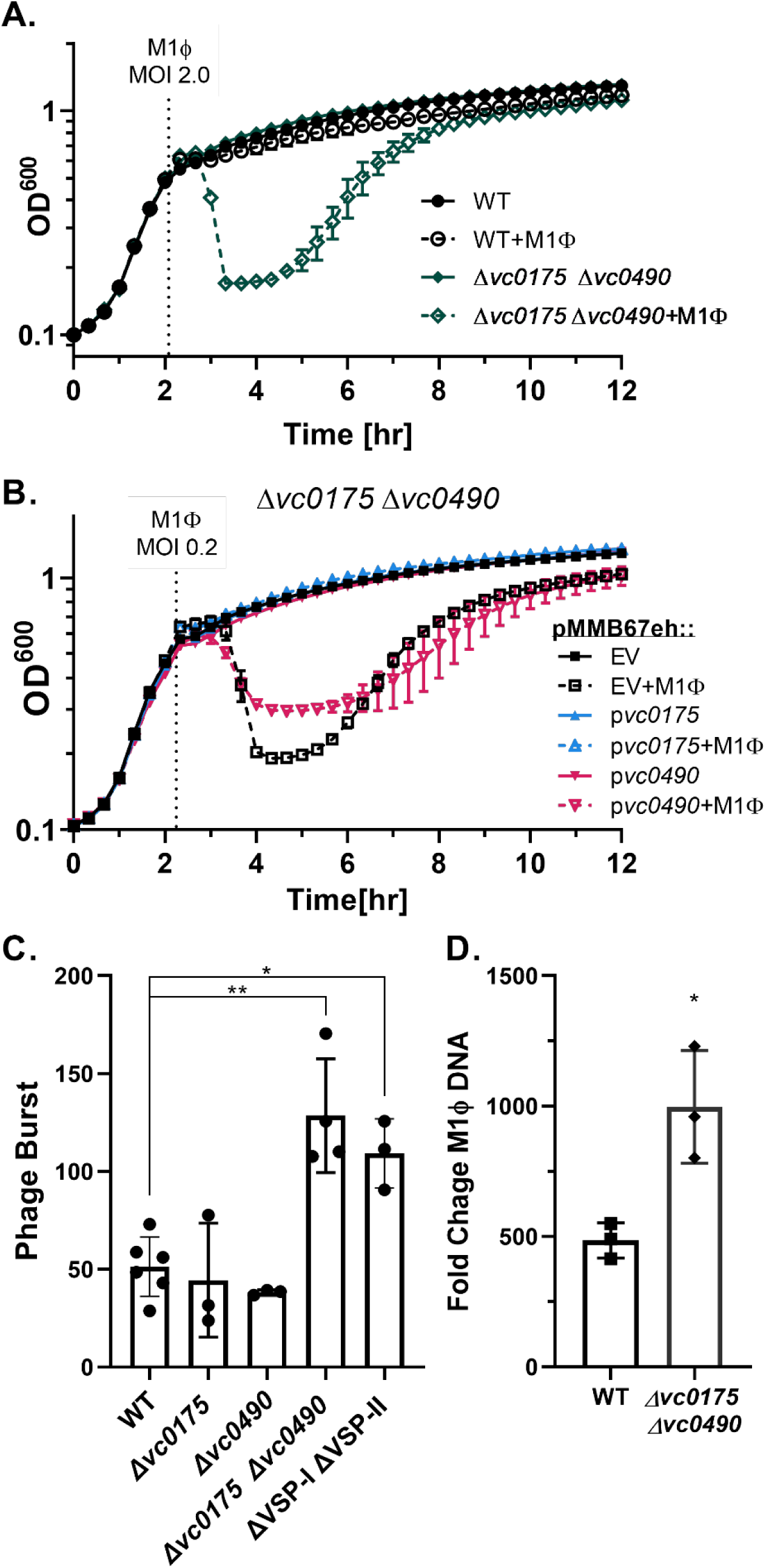
*vc0175 and vc0490* protect growing cultures of *V. cholerae* from M1Φ infection. (A) Growth and lysis curves of *V. cholerae.* Indicated strains were infected with M1Φ at an MOI of 2.0 at the time displayed by the dotted line. All strains were grown in LB shaking at 37°C and optical density was read at regular intervals with a plate reader. (B) *V. cholerae* Δ*vc0175* Δ*vc0490* strain containing the plasmid pMMB67eh or the same plasmid expressing *vc0175* or *vc0490* was grown and infected with M1Φ at an MOI of 0.2. (C) Phage burst assays. Strains were grown to mid-log phase before being infected with phage at an MOI of 0.1. After a period of absorption cultures were further diluted 1/2500,1/25000, and 1/250000. Burst is calculated by dividing the total output phage by absorbed phage (T0-T10). Strains are indicated on the x-axis. P-values were determined by Unpaired T-tests with Welch’s correction and significance shown as *p<0.0332 and **p<0.0021. (D) Fold change in M1Φ genome copy through infection. The indicated strain was grown to mid-log phase before being infected with M1Φ at an MOI of 0.02 and grown in LB at 37°C on a roller. Immediately upon infection and at 20 minutes post infection a portion of the culture was removed and boiled. The boiled culture was then diluted 1:50 and used as template for qPCR with primers targeting the ICP3 DNA polymerase. Fold change was calculated by dividing the CT value at 20minutes by the value at 0 minutes for each culture.

We were able to restore the protection by expressing these genes individually from a plasmid in the Δ*vc0175* Δ*vc0490* mutant. While expression of either gene from a plasmid reduced the overall lysis of a culture, *vc0175* had a stronger effect in the conditions tested, nearly entirely ablating lysis (**Fig. 3B**). Ectopic expression of *vc0490* did not completely prevent lysis but reduced the amount a culture lysed significantly versus an empty vector (**Fig. 3B**). This further indicates that that either gene is sufficient for protection against M1Φ. The lack of toxicity to the host when these genes are overexpressed also suggests that they are not inhibiting phage replication by simply hindering the *V*. *cholerae*’s ability to grow **(Supplemental Fig 2A).** We did however see a mild growth defect when the genes were expressed in *E. coli,* indicating that these genes cause minor growth inhibition in other bacteria **(Supplemental Fig. 2B).** Although to a smaller extent, protection to phage-induced cell lysis was still observed at a higher MOI in the Δ*vc0175* Δ*vc0490* mutant when either gene was over-expressed **(Supplemental Fig. 2C).**

The sufficiency of either gene to protect against productive M1Φ infection is also shown via a burst assay. While M1Φ produced approximately 50 phage per cell in VSP^+^ (WT), Δ*vc0175,* and Δ*vc0490* strains (**Fig. 3C**) the burst size was doubled in strains with either both VSPs or both *vc0490* and *vc0175* deleted, resulting in ∼100 phages released per cell (**Fig. 3C**). This increase in burst could help partially explain the larger plaque size observed in these strains (**Fig. 2A**) although it may not be the only contributing factor [31][32]. The burst sizes of M1Φ on these mutants that is nearly 2.5 times of WT which also matches the EOP data closely (**Fig. 2A**, **Fig. 3C**).

Next, we tested if viral DNA replication of M1Φ was inhibited by *vc0175* and *vc0490* in *V. cholerae*. To do this, we measured by qPCR the increase in viral DNA over the course of a replication cycle. As predicted, we observed more viral DNA in the Δ*vc0175* Δ*vc0490* mutant than the WT. This difference of ∼2x more DNA produced closely matched the overall burst differences (**Fig. 3D**). These results suggest that VC0175 and VC0490 are most likely targeting phage DNA replication, reducing the overall number of phage genomes produced in each cell, and thereby reducing the total number of phage released from each cell.

VC0175 (AvcD) has been shown to be a functional cytidine deaminase capable of converting dCMP to dUMP, presumably changing the nucleotide pool in the cell to inhibit phage replication [19]. Another study showed that VC0490, working together with VC0491 and VC0492, decreased plasmid stability in El Tor biotype by an unknown mechanism [24]. Similar to the previous study, we observed that VC0490 appeared to be a SMC-Like protein based on Alphafold predictions [33]. These proteins are often required for proper chromosome segregation during replication and can play a role in plasmid maintenance.

Although VC0175 and VC0490 both target phage replication, because of the dissimilar predicted functions of these two systems, we reasoned that each system could have a unique target specificity. To evaluate if both systems similarly reduced plasmid uptake and stability, we measured the efficiency of strains to take up commonly used plasmids via conjugation from *E. coli.* Significantly more transconjugants were obtained in the Δ*vc0490* recipient than those from WT or the Δ*vc0175* mutant (**Fig. 4**). This phenotypic divergence suggests that these two systems target MGEs with different specificity and it is probable that these systems use two distinct mechanisms to inhibit M1Φ DNA replication.

**Figure 4:**
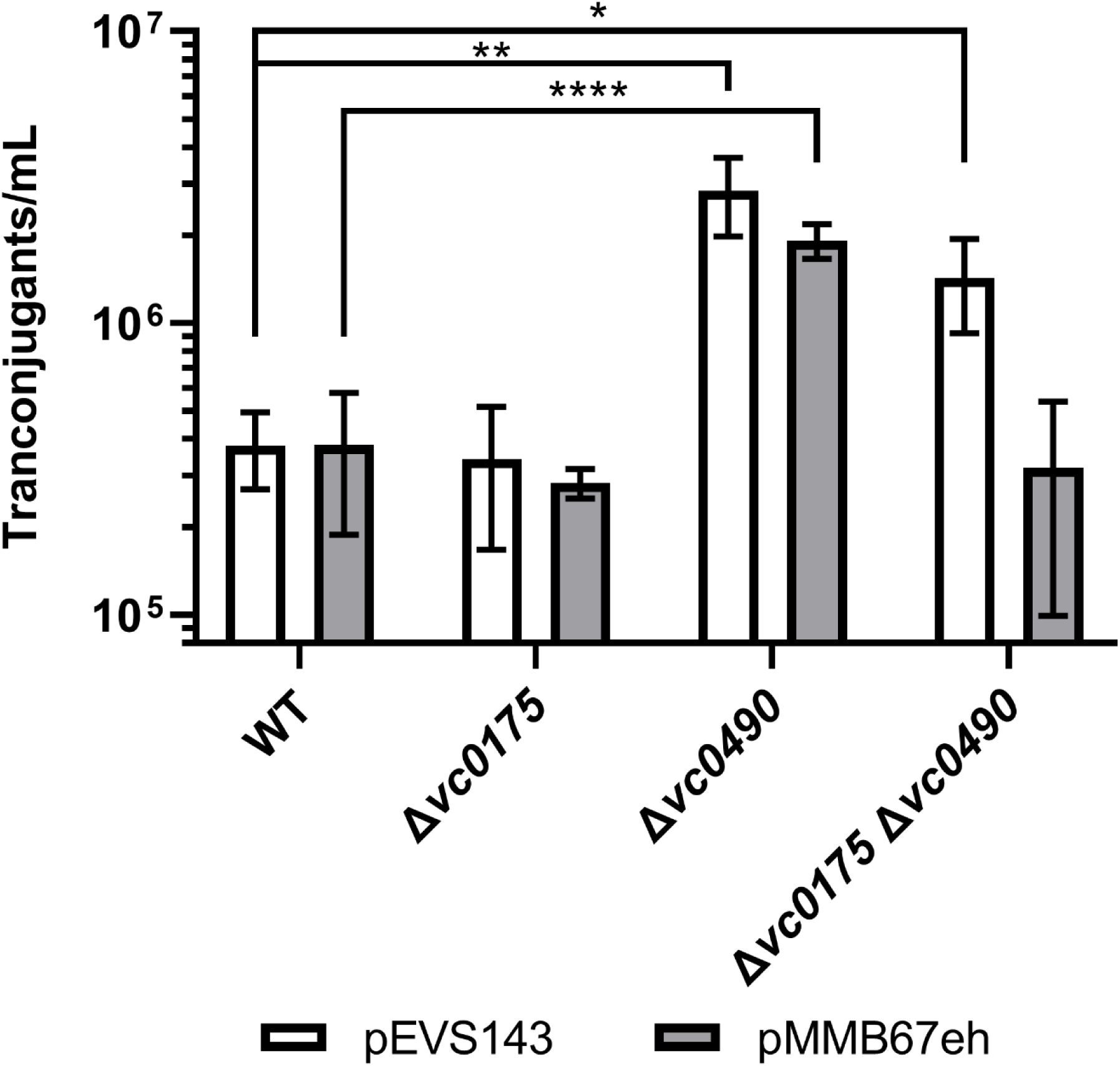
VC0490 reduces plasmid conjugation. *E. coli* donors carrying either pEVS143(white bars) or pMMB67eh (grey bars) were mixed with different *V. cholerae* recipients as indicated and incubated together for 2.5 hours. Cells were then resuspended and an aliquot was plated on LB containing polymyxin B which is lethal to *E. coli* and either kanamycin (for pEVS143) or ampicillin (for pMMB67eh) and incubated overnight before the resultant colonies were counted. Significance was determined by Unpaired T tests with Welch’s correction with significance shown as *p<0.0332 and **p<0.0021 ****p<0.0001.

### A SNP in ICP3_2016_M1 results in VSP sensitivity

Previously identified ICP3 did not show this susceptibility to the VSPs. Therefore, we sought to determine what causes M1Φ to be susceptible to the defense afforded by these two genes. As previously stated, M1Φ has 99.5% identity and 99% coverage when compared to a previously sequenced ICP3_2009_A. While this was sufficient for us to categorize the isolated phage as ICP3, hundreds of differences remained between these two related phages. To isolate VSP resistant ICP3 with a more similar genetic background to M1Φ, we used the same RWS sample where M1Φ was isolated, and we re-isolated another ICP3 phage using the same scheme but for phages that did not show a preference for hosts with or without the VSPs. We successfully isolated a naturally occurring VSP resistant phage that we called ICP3_2016_ M2 (M2Φ). Sequencing of this phage revealed that its genome was almost identical to M1Φ, with only 2 single nucleotide polymorphisms (SNPs) (Accession number ON464736). Both of these SNPs map to the gene that encodes the well conserved phage polymerase gp22 (**Supplemental Fig. 3)**. These VSP sensitivity determining mutations are both found in the exonuclease domain of the phage DNA polymerase but only one would result in an amino acid change (L94I). While this change is subtle, it occurs in a well conserved region proximal to the invariant non-catalytic aspartate that is essential for the ssDNA exonuclease function of the polymerase [34].

The minor genetic differences in M2Φ appeared to have a substantial impact on its ability to tolerate *vc0175* and *vc0490* as there was no change in EOP when comparing plaques formed on the mutant strain to WT (**Fig. 5A**). This is in direct contrast to the increase in EOP we observed on the same strains with M1Φ (**Fig. 2B**). When lysis assays were performed with each phage infecting at the same MOI (**Fig. 5B**), WT resisted lysis by M1Φ but not with M2Φ (**Fig. 3A and 5B**). It is also notable that while both M1Φ and M2Φ phages lysed the ΔVSP host strains to a similar degree, M2Φ appeared to initiate host cell lysis earlier than M1Φ (∼20 mins for M1Φ vs ∼40 mins for M2Φ after phage addition), potentially suggesting differences in replication between the two phage. Our results suggest that M2Φ is not sensitive to inhibition conferred by *vc0175* and *vc0490;* and we predict the resistance is likely due to a change in the structure and/or function of the phage DNA polymerase.

**Figure 5:**
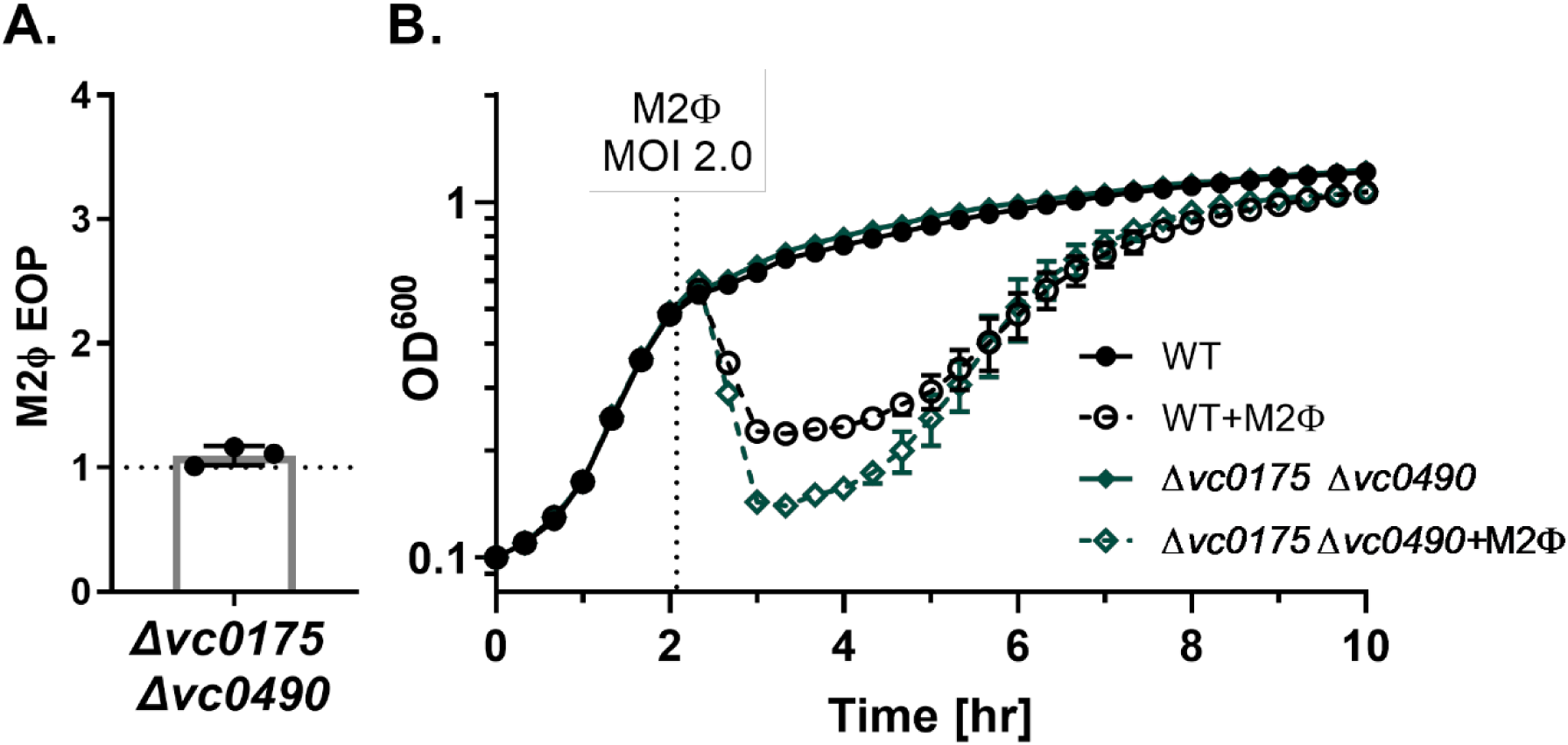
M2Φ is not susceptible to VC0175 and VC0490. (A) EOP of M2Φ on Δ*vc0175* Δ*vc0490* compared to the parental VSP^+^ strain (WT). (B) Growth and lysis curves of the indicated strains with or without M2Φ added at an MOI of 2.0 at the time shown with a dotted line. Uninfected data is the same as shown in Figure 3A.

### Protection by *vc0492-490* is quorum sensing dependent

Phage infection relies on the active metabolism of the cell and therefore it is expected that many phages replicate more efficiently while strains are actively growing in mid-log phase. However, when optimizing infection conditions, we noticed a striking difference in lysis when phages were introduced at different cell densities of the *V. cholerae* host. During our standard infections with M1Φ at mid-log growth phase (OD ∼0.5), we observed the expected *vc0175 vc0490* dependent protection (**Fig. 6A**). Expectedly, at higher cell density (OD ∼0.8), no lysis was detected at all, which we assumed was due to the reduced metabolic state of the host at late log phase that was not conducive to phage infection (**Fig. 6A**). Intriguingly, at low cell density (OD ∼0.2), we observed lysis in all strains infected with M1Φ (**Fig. 6A**), which is in stark contrast to what we would have expected from the phage defense elements on VSP-I or VSP-II. These results suggest that the VSP phage defense systems are not active at low cell density and the phage protection phenotype at a higher cell density (OD ∼0.5) is a sign of more complex regulation.

**Figure 6:**
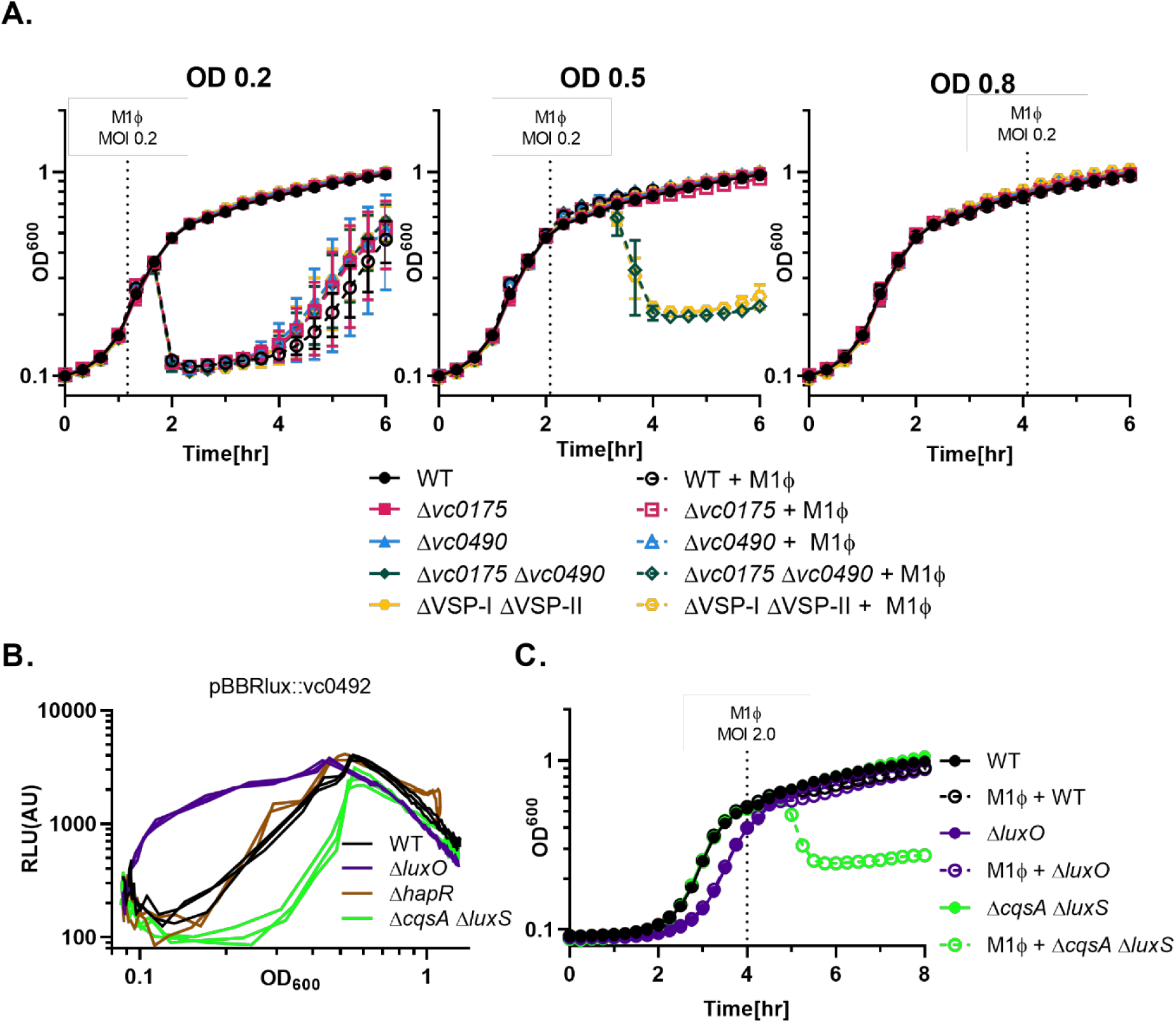
Defense by VC0490 is controlled by quorum sensing. (A) Growth and lysis curves of *V. cholerae* infected by M1Φ at different cell densities. M1Φ was added at an MOI 0.2 at the time indicated with a dotted line that corresponds closely with the OD_600_ shown at the top of each graph. Strains used for all three graphs are indicated on the far right of the figure. (B) Quorum sensing mutant strains of *V. cholerae* carrying the reporter plasmid pBBRlux with the *lux* operon under the control of the region upstream of *vc0492* were grown at 37°C with shaking. Optical density (OD_600_) and bioluminescence were measured at regular intervals. Relative luminescence (RLU) was calculated by dividing the luminescence by optical density and was then plotted relative to the OD_600_ to show the normalized bioluminescence throughout growth. (C) Grow and lysis curves of C6706 (WT) and various quorum sensing mutants upon M1Φ infection. M1Φ was added to the indicated strains at the dotted line at an MOI of 2.0.

While *vc0175* was previously shown to be important for quorum-sensing (QS) dependent multicellular aggregate formation in *V. cholerae* at high cell density [35], the link between *vc0490* and QS is less clear. To test if *vc0490* is also regulated by QS, we used a reporter plasmid containing the *Photorhabdus luxCDABE* luciferase operon under the control of the region immediately upstream of *vc0492* to measure gene expression in different QS mutants. In the Δ*luxO* mutant that is genetically locked at QS high cell density state [36], there was an increase in bioluminescence across different cell densities when compared to the WT. In contrast, in the

Δ*cqsA* Δ*luxS* strain missing both the two known QS autoinducer synthases, and thus locked at low cell density QS state [36], there was a decrease in bioluminescence across different cell densities when compared to the WT. Unlike the Δ*cqsA* Δ*luxS* strain, a *hapR* mutant which is also locked at low cell density QS state [37], did not show any change in luminescence. Together, these data strongly suggest that the *vc0492-490* operon is controlled by QS [38] and is active only at higher cell densities. Specifically, the inactivity of this particular system at low cell density is due to the repression of gene expression by the Qrr sRNAs rather than the lack of a transcriptional activation by HapR, a mechanism that has been previously observed [39, 40]

To test the QS-dependent phage protection, we infected different QS mutants with M1Φ at OD ∼0.5 where we would expect both *vc0175* and *vc0492-490* to be active and protective. Indeed, we observed complete protection in the WT strain as well as the Δ*luxO* mutant even at a high MOI, but not in the Δ*cqsA* Δ*luxS* strain. These results strongly suggest a functional connection between QS and protection from M1Φ infection.

## DISCUSSION

In this work, we have isolated a circulating novel variant of the *Vibrio cholerae* phage ICP3 (M1Φ) in a cholera patient stool sample that is susceptible to defense genes carried on the *Vibrio* Seventh Pandemic (VSP) Islands. We determined that both *vc0175* on VSP-I and *vc0490* (or the *vc0490-2* operon) on VSP-II are anti-phage elements that target M1Φ. We show that these systems inhibit DNA replication of the phage, reducing the overall number of phage released per cell. Moreover, these systems significantly protect *V. cholerae* cultures from lysis by M1Φ at both high and low MOIs. We have also isolated in the same stool sample a naturally occurring phage that is closely related to M1Φ that is VSP-resistant (called M2Φ), carrying changes in the proofreading domain of the phage DNA polymerase.

As previously stated, there are many SNPs throughout the genomes which differentiate the previously isolated ICP3 and the newly isolated ICP3_2016_M1 Φ or M2 Φ including the gene that encode GP22 (DNA polymerase). In contrast, there is exactly one predicted amino acid divergence in GP22 for the two new phage isolates **(Supplemental Fig. 3)**. Among the related T7 polymerase family, this residue is mostly a leucine and is next to the well conserved non-catalytic aspartate [34]. While M1 has a leucine in this position, M2Φ has an isoleucine. Looking through deposited sequences via the Basic Local Alignment Search Tool (BLAST), we were unable to identify any ICP3 or T7 sequences that had a similar leucine to isoleucine change at the equivalent position in the DNA polymerase. Therefore, M1Φ appears to be the ancestral isolate and it is very likely M2Φ is derived from M1Φ to allow better replication in circulating VSP- containing *V. cholerae*. However, the change in the M2Φ DNA polymerase may come with a fitness cost due to the high conservation of this region which suggests functional importance.

During the course of this study, three independent studies [19,20,24] identified the same gene clusters on VSP-I and VSP-II as potential anti-phage systems. Our work complements these findings but, importantly, shows function of these two islands in phage defense in their native host species. By identifying both sensitive and resistant *V. cholerae* phage isolates from the same patient stool sample, we have directly observed the arms race between the phage and its native bacterial host, illustrating how phage counteracts these defense systems within the native environment (i.e., within the human small intestine) and providing additional insights on the molecular mechanisms used by these systems for defending against phage attack.

VC0175 (also known as DcdV or AvcD [19]) was identified to be a cytidine deaminase that functions to modify nucleotide pools. Interestingly, a small RNA called AvcI upstream of *vc0175* controls the activity of VC0175 post-translationally, resembling a toxin-antitoxin system [41]. Over- production of VC0175 in the absence of AvcI leads to cell filamentation, a phenomenon similar to thymine-less death (TLD), even though VC0175 only carries out deamination of both dCMP and dCTP. Expression of VC0175 and its homologs from other species in *E. coli* leads to toxicity but also protection against some coli phages, suggesting modification of the nucleotide pools inside the bacterial host could lead to phage defense via direct inhibition of phage replication [19, 20]. Our study uniquely shows that M1Φ is sensitive to alteration of the nucleotide pools mediated by VC0175 in a system with no observed toxicity. This idea was further supported by our isolation of the VSP-insensitive M2Φ phage variant in which a polymorphism was identified in the exonuclease domain of the phage DNA polymerase. We predict that the M1Φ phage DNA polymerase is more sensitive to an imbalanced nucleotide pool, and this sensitivity could lead to blockage of replication or an “error catastrophe” that reduces phage fitness.

Deletion of *vc0490* or the whole *vc0492-0490* operon (also known as *ddmABC*) mimics the effect of removing VSP-II for phage defense. Recent study suggests that the three gene products function together to reduce plasmid stability in the pandemic *V. cholerae* El Tor biotype [24]. This system appears to be especially active towards small or medium sized plasmids [24]. Our data further support that *vc0492-490* reduces conjugation of certain mid-sized plasmids (4.3kb and 8.8kb) from *E. coli* to *V. cholerae*. Expression of this operon in *E. coli* also confers resistance to some coliphages [24]. Our study further illustrates the importance of this operon in defending against a native circulating phage. The exact mechanism for phage protection conferred by this system remains unclear. Using bioinformatic analysis and structural prediction using Alphafold, we and others [24] predicted that VC0490 contains a SMC domain common in certain chromosome partition proteins and therefore it is likely needed for recognizing and binding to specific features in the target DNA. VC0492 is predicted to be an endonuclease, likely for processing and degrading target DNA. The small VC0491 protein is predicted to be an ATPase which could function with nearby effectors to respond to foreign genetic materials [42]. It is therefore most likely that the nuclease activity of VC0492 is being directed and regulated by VC0490 and VC0491. Yet, the specific target and activator of this system remain elusive. VSP- resistant phage M2Φ, producing a slightly different DNA polymerase than that from the VSP- sensitive M1Φ, provides a clue that this defense system could either sense the polymerase directly, or more likely the phage DNA replication intermediates associated with the polymerase. Indeed, a phage defense system called Nhi, carrying both a nuclease and helicase domain, is recently identified to sense phage replication intermediates to direct DNA processing [43]. Regardless of the mechanism, although many versions of VSP-II circulate in the natural isolates of *V. cholerae*, nearly all VSP-II islands contain the *vc0492-490* operon [21, 44], suggesting there is significant pressure to maintain these core genes.

Moreover, recent study on VC0490-492 predicted the mechanism of defense to be abortive infection (ABI) [24], where a small number of infected bacterial cells sacrifice themselves to halt the propagation of the phage and protect the rest of the uninfected cells in the same population [45]. Data from our study suggests that ABI, at least in *V. cholerae*, may not be the sole acting mechanism for these systems. *V. cholerae* carrying functional VSPs appears to be fully protected from M1Φ at both low MOI of 0.2 (**Fig. 6A**) and high MOI 2.0 (**Fig. 3A**), suggesting that infected cells are still viable when phage are abundant. A defense system using ABI would normally display characteristic cell death at high MOI and not result in a productive infection at low MOI as we have observed. Lack of abortive infection at high MOI implies that a specific molecule is being targeted/sensed by the defense system and is usually absent or present infrequently in the bacterial host. Since VC0490-2 (or DdmABC) act on a variety of plasmids as well as phages, the target likely is not a specific protein encoded by these foreign genetic elements. Again, based on the polymorphism existing between M1Φ and M2Φ, we speculate some replication intermediates are targeted or sensed by this system. Further analyses are required to identify the exact molecular mechanisms, but our work provides a phage-host combination that readily allows for such analysis.

While protection from phage through heterologous expression in foreign host such as *E. coli* is a very powerful approach, especially in large scale studies where demonstration of the phage defense activity is the key; these approaches may not inform the intricate functions of some anti-phage systems. Our study uniquely illustrates that complex gene regulation exists for these phage defense genes in their native host environment and suggests conditions where phage defense systems are coordinated with other cellular processes. For instance, we show that these defense systems are under both growth phase and quorum sensing control: these systems are only expressed and become active at certain cell density, and therefore M1Φ phage cannot infect and lyse *V. cholerae* host carrying these defense systems at high cell density (e.g., late exponential phase/early stationary phase) unless quorum sensing (QS) is disrupted. Dependence of QS for phage protection in *V. cholerae* has been reported previously, however, such protection is mediated through a HapR-dependent production of haemaglutinin protease, and partly through downregulation of phage receptors [46]. In contrast, the QS-dependent protection we identified in this study is HapR independent. What is the possible driving force for connecting QS and phage protection? Since *V. cholerae* only develops natural competence and picks up foreign DNA at high cell density mediated by QS [47], QS mediated defense against foreign DNA could act as a checkpoint and prevent acquisition of unwanted MGEs by activating these defense systems.

While our ability to identify anti-phage systems has expanded rapidly, our data suggests that there are regulatory and mechanistic insights that can only be gained by studying these systems in their native hosts defending against endogenous phage. Outside of *V. cholerae,* there has also been increasing awareness of the arsenal of anti-phage genes contained on MGEs [48, 49] and many components from these systems have been used as basic molecular biology tools for years [50, 51]. As phages are so plentiful and diverse, undoubtedly there are more anti-phage mechanisms that are yet to be uncovered. Indeed, many (non-temperate phage) MGEs themselves are thought to be derived directly from phages, or derelict phage themselves [52, 53]. By identifying new systems on these abundant but often ill-defined islands, we can help expand the “molecular toolbox” and pave the way for future innovation.

## ACKNOWLEDGEMENT

We thank Dr. Andrew Camilli and Dr. Chris Waters for providing some bacterial strains and plasmids for use in this study. We also thank them and their lab members for insightful comments and discussions. WLN and BJO were supported by R01AI121337, and BJO was supported by T32GM007310. MA thanks the icddr,b hospital and laboratory staffs for their support, and the Governments of Bangladesh, Canada, Sweden, and the United Kingdom for core/unrestricted support.

## MATERIALS AND METHODS

### Bacterial strains and Growth Conditions

All *V. cholerae* strains used in this study were O1 El Tor. Bacterial strains utilized in this study are listed in **Supplementary Table 2**. WN6145 was the primary *V. cholerae* strain used in the study. This strain is derived from E7946, an El Tor Ogawa strain [54] to promote phage infection. Genetic alterations include elimination of phase variable site in the gene that encodes the common phage receptor the O1-antigen [55], deletion of the CTX prophage, and deletion of the kappa prophage. Bacteria were propagated in LB at 37°C shaking with aeration. Strains containing pMMB67eh were grown with 100µg/mL Ampicillin, strains with pBBRlux::*vc0492* were grown with the addition of 2.5µg/mL Chloramphenicol. For growth curves, strains were grown in 200µL cultures in 96-well plates (ThermoScientific Nunc 167008) at 37°C with lid shaking.

### Generating *V. cholerae* Mutant Strains

Natural transformation was used to introduce antibiotic resistance markers into *V. cholerae* [56]. Unmarked deletions for the desired gene or gene cluster were made by splicing by overlap extension (SOE)-PCR and the resulting PCR products was transformed at the same time with PCR products where a cassette conferring kanamycin or spectinomycin resistance replacing *lacZ.* After selecting on the appropriate antibiotic colonies were screened for deletion of the unmarked locus [57]. Strains were made competent by growth on sterile chitin in instant ocean [56]. For marked constructs, approximately 500ng of PCR product was added to competent cells on chitin. When co-transforming, 200ng DNA containing the antibiotic resistance marker was used with 2µg unmarked PCR product. The oligonucleotide sequences used for PCR and sequencing reactions will be provided upon request. Plasmids were constructed via digestion with restriction endonucleases and subsequent ligation into vectors with T4 ligase. Insertions were initially identified by PCR and confirmed via sanger sequencing.

### Isolation of *Vibriophage* from patient stool samples

Stool samples from cholera patients were previously collected as described [58]. Phage were isolated by scraping a small amount of -80°C stool sample into a 1.5mL Eppendorf tube. These were thawed and gently spun to remove bacteria and other debris before the supernatant was passed through a 0.45µM filter (VWR 28145-485). Soft agar overlays were prepared by growing ΔVSP*-*I ΔVSP-II (WN7006) to OD ∼0.5 at 37°C shaking, adding this culture to 0.5% Top Agar, and overlaying this mixture on LB plates. Serial dilutions of filtered stool samples were then applied on top of the overlay and incubated at 37°C overnight. Single plaques were picked into 25µL TM buffer. Five µL of the plaque suspension was boiled, diluted 1:20 and used as template for ICP phage identification PCR using previously described primers [5]. Serial dilutions on these picked plaques were performed and plated on soft agar overlays with either ΔVSP*-*I ΔVSP-II (WN7006) or the isogenic VSP^+^ strain (WN6145) *V. cholerae* to assess differences in plaquing between the two hosts. For those phages that were further studied, single plaques were picked from the ΔVSP*-*I ΔVSP-II (WN7006) plates of this second plating to subsequently have phage stocks prepared via PEG precipitation.

### Phage infection

Phage stocks were generated on the permissive ΔVSP*-*I ΔVSP-II strain (WN7006) using polyethylene glycol (PEG) precipitation from liquid cultures. *V. cholerae* was grown to OD_600_ ∼0.3 in LB at 37°C shaking, phage was added at an MOI of 0.2. Cultures were monitored until fully lysed (cleared). At this point DNAse, RNAse, and 0.002% (v/v) chloroform was added to eliminate bacteria and bacterial DNA/RNA. Bacterial debris were removed by adding NaCl to a concentration of 0.5M and centrifuging at 4°C. The supernatant was then taken and 10% PEG 8000 was added. After overnight precipitation cultures were spun and the resultant pellet of phage was resuspended in TM buffer (10mM Tris-HCL pH 7.5, 100mM NaCl, 10mM MgCl_2_). Phage stocks were further cleared by the addition of chloroform once more and centrifugation. The aqueous phase from this final centrifugation was then used as a working phage stock. Titering of phage for was carried out by growing *V. cholerae* to mid-log, infecting cultures with diluted phage stocks, and allowing absorption to occur for 5-10 minutes before plating on 0.7% top agar. After a brief amount of time to solidify, plates were incubated at 37°C overnight. Plaques were enumerated with the aid of magnification to ensure count of small plaques. EOP was calculated after titering equal number of phage on different *V. cholerae* strains at the same time ensuring that phage titer and incubation times remain the same. MOI was determined by calculating the colony forming units (CFU) per milliliter at a given OD_600_ value and then adjusting the number of plaque forming units (PFU) to be added based on the desired MOI and expected CFU. Average burst size was determine by using one-step growth curves[59] performed with at least three biological replicates. Phage burst is reported as individual values with a bar representing the median ±SD (Standard Deviation) in Fig. 2B. Bacterial growth and lysis curves of *V. cholerae* with ICP3_2016_M1 (Fig. 2 & 6 Supplementary Fig. 2) or M2 (Fig. 5) were performed with phage added at the indicated time and MOI at 37°C shaking. For complementation experiments using pMMB67eh, LB broth and top agar was supplemented with 100µg/mL Ampicillin and 100µg/mL IPTG (isopropyl-β-D- thiogalactopyranoside).

### Quantification of plaque size

After plaque plates had been incubated and counted plaque size was evaluated. To increase contrast of the plaques an overlay of 0.1% (2,3,5-Triphenyl-2H-tetrazolium chloride(TTC) (Thermofisher AAA1087009) was applied [60]. Plates were imaged on an Epson Perfection V800 Dual Lens scanner. Analysis of the images was performed in the program Fiji. Images were smoothed, the “Find Edges” function was used before automatic Thresholding (Li). The “analyze particles” function was then used to identify plaques selecting those larger than 4 pixels and with a circularity between 0.8 and 1.0.

### Isolation of Phage DNA and sequencing”

Phage gDNA was isolated from lysate. Lysate was treated with DNase and RNase to remove bacterial DNA at 37°C for 30 minutes before heat inactivation at 65°C for 10 minutes. Phage capsids were disrupted by the subsequent addition of proteinase K and SDS. Phage DNA was then cleaned up using Zymo DNA Clean & Concentrator kit (D4029) and eluted in water. Illumina libraries were prepared for each phage using the standard Nextera XT DNA Library prep protocol. Sequencing was conducted at the Tufts University Core facility using an Illumina NextSeq 550 and Single-end 75nt reads. Genome sequence analysis was performed with CLC Genomic Workbench v20.0.4.

### Real-time quantitative PCR

Reactions for qPCR experiments were carried out with PowerUp SYBR Green Master Mix (Thermofisher) on a CFX Connect Real-Time PCR Detection System (Bio-Rad). Three independent samples were tested for each strain and each template was assayed in technical duplicate. Bacteria were grown to OD=0.5 at 37°C with aeration, at which point phage were added at an MOI = 0.1. Immediately upon infection and 20minutes post infection 20µL samples were taken, boiled, and diluted 1:50. This boiled lysate was used as template for qPCR using phage specific primers as previously described [5] with primers WNTP1342 (5’- ATTGTCGAGTGGGACAAAGG-3’) and WNTP1343 (5’-ACCAACTCGACGCATAGCTT-3’).

### Conjugation

Strains were grown overnight in LB at 37°C shaking. *V. cholerae* strains used for these experiments contained Δ*lacZ::*spec (WT WLN7019, Δ*vc0175* WLN7008, Δ*vc0490* WLN7011, and Δ*vc0175* Δ*vc0490* WLN7020) for enhanced selection of transconjugants. Equal quantities of *E. coli* donor and *V. cholerae* recipient were mixed and spun gently before being resuspended in 80µL LB. This concentrated mixed was plated in 3x 20µL spots on LB with no drug supplements and incubated at 37°C for 2.5 hours. Bacteria was scraped from the plates and resuspended in LB before being serially diluted and plated on LB polymyxin B (50U/mL) Spectinomycin (100µg/mL) and either Ampicillin (100µg/mL) for pMMB67eh or Kanamycin (100µg/mL). Colonies that arose after overnight incubation at 37°C were enumerated.

---

**Supplemental Figure 1:**
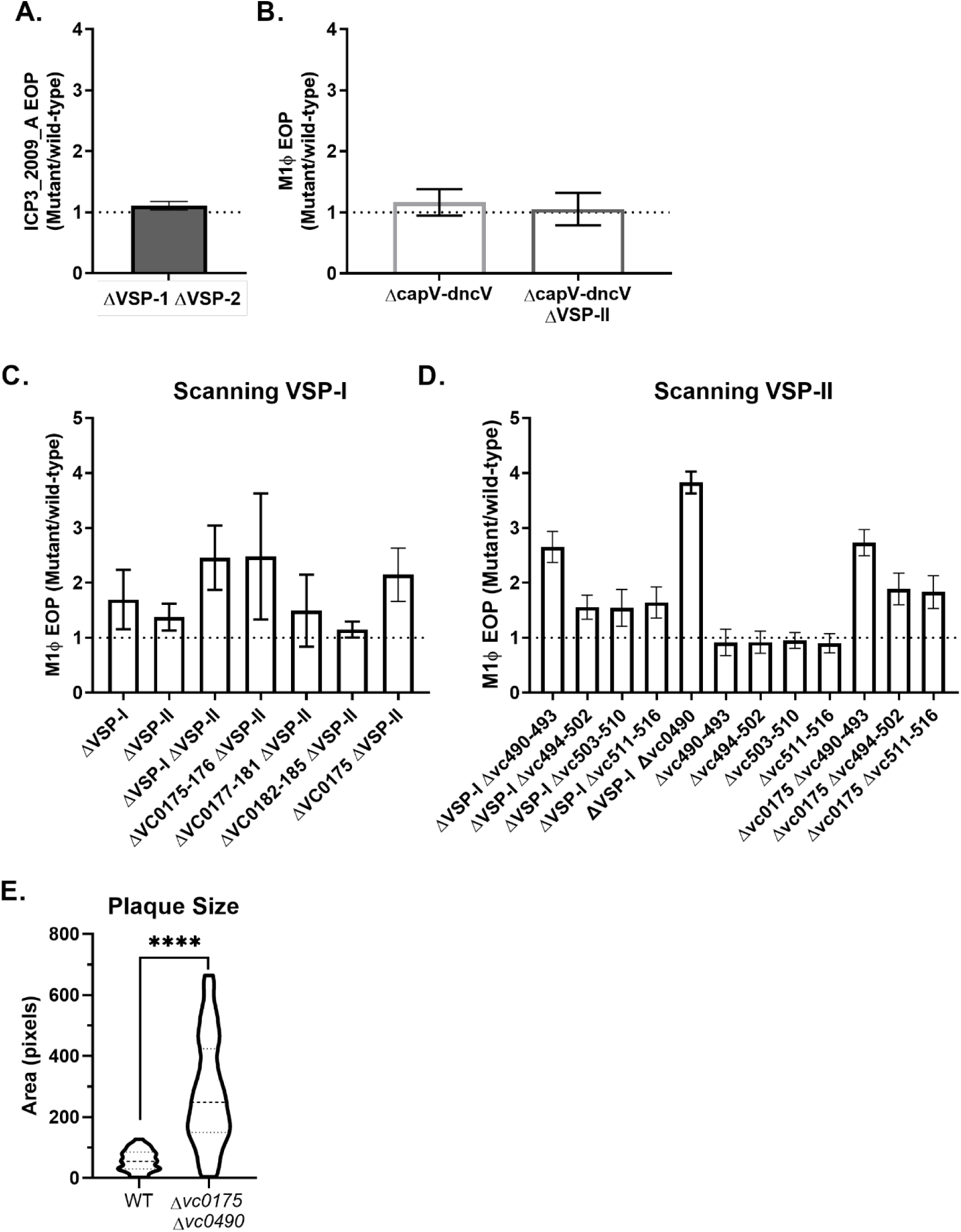
Determining the anti-phage elements on VSP-I & VSP-II. (A) EOP of ICP3_2009_A infection determined on the *V. cholerae* ΔVSP-I ΔVSP-II mutant and the parental VSP^+^ strain (WT). (B) EOP of ICP3_2016_M1 (M1Φ) infection determined on the *capV-dncV* mutant and WT. (C) EOP of M1Φ infection determined on different VSP-I related mutants and WT. Values from ΔVSP-I, ΔVSP-II, and ΔVSP-I VSP-II are also shown in Figure 2 and repeated here for clarity. (D) EOP of M1Φ infection determined on different VSP-II related mutants and WT. (E) Quantification of plaque sizes. Plaques were imaged and processed with Fiji to determine the area of each plaque on a plate. Sizes are from plates from two separate days. Significance was determined by Kolmogorov–Smirnov test where ****=p<0.0001.

**Supplemental Figure 2:**
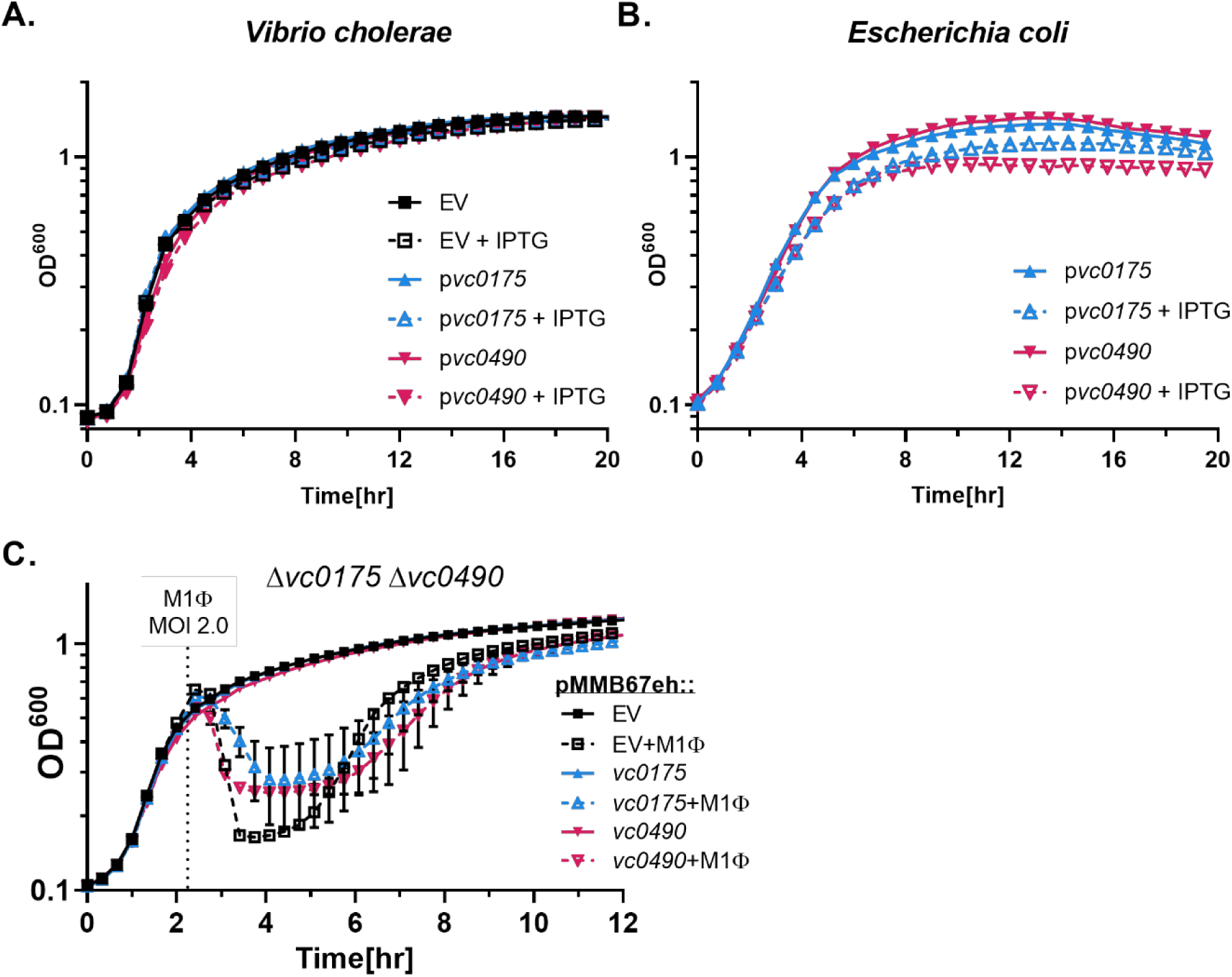
Effect of vc0175 and vc0490 expression on bacterial growth. Growth curves of *V. cholerae* (A) and *E. coli* (B) carrying the plasmid pMMB67eh with the indicated insert +/- 100µM IPTG. Average of at least three biological replicates are shown. (C) *V. cholerae* Δ*vc0175* Δ*vc0490* strain containing the plasmid pMMB67eh or the same plasmid expressing *vc0175* or *vc0490* was grown and infected with M1Φ at an MOI of 2.

**Supplemental Figure 3:**
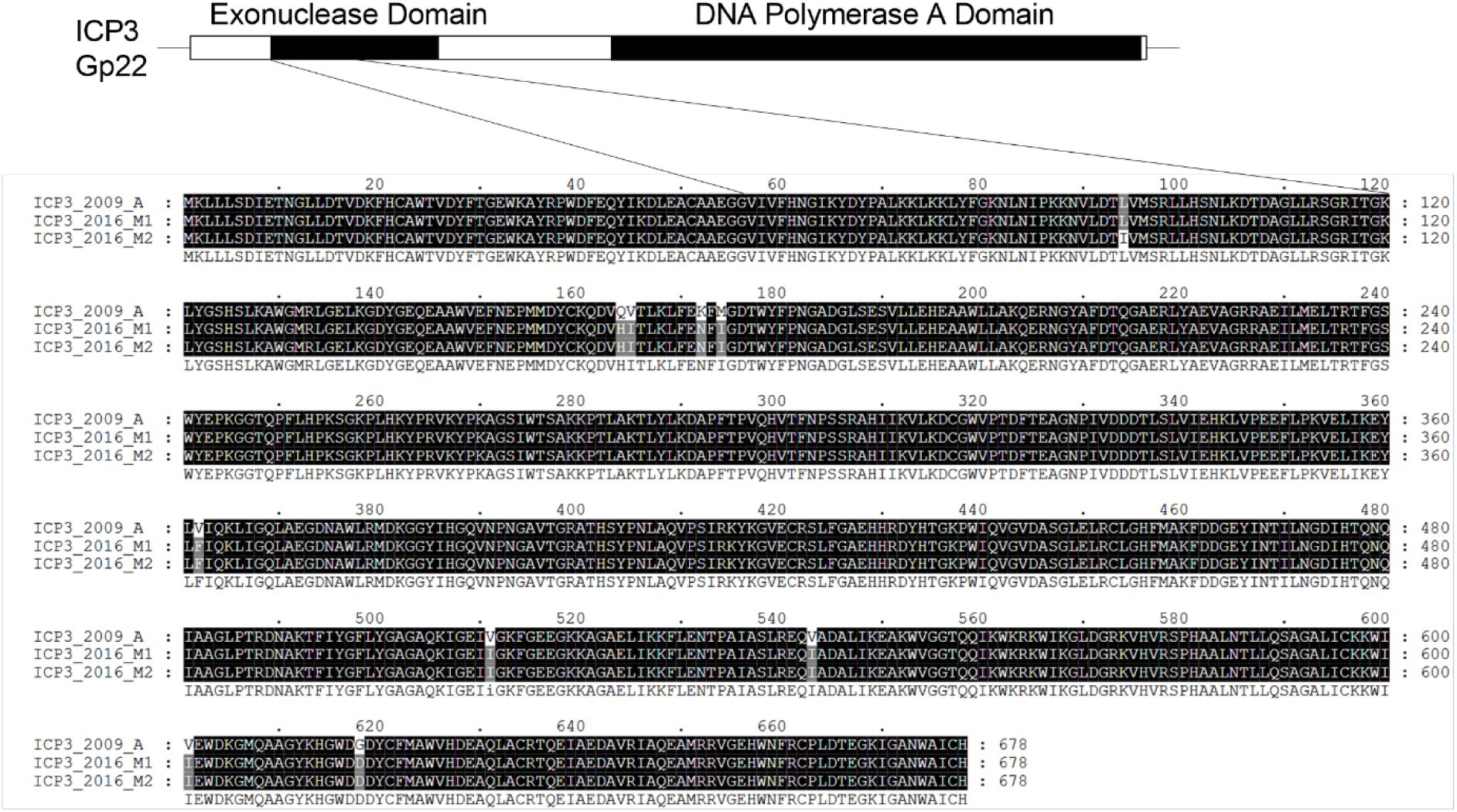
Polymorphism in the ICP3 DNA Polymerase (GP22) Scale diagram of the ICP3 polymerase GP22 with the black boxes indicating the well conserved exonuclease and DNA Polymerase A domains. Breakout alignment of M1 and M2 gp22 was performed with T-Coffee and display was generated using GENEDOC. Single amino acid change can be seen as black text on a grey background.

**Supplemental Table 1:** Summary of phage found in stool samples

|  | ICP1 | ICP2 | ICP3 | No Bands |
| --- | --- | --- | --- | --- |
| Sample 1 | ND | ND | ND | 0% |
| Sample 2 | 94% | ND | ND | 6% |
| Sample 3 | 13% | 13% | 38% | 38% |
| Sample 4 | ND | ND | ND | 0% |

Picked plaques were boiled, diluted, and used as template for diagnostic PCRs using primers targeting the conserved polymerases of ICP1, ICP2, and ICP3. Percent of plaques with indicated band shown. No Bands indicates no bands were detected after two separate attempts to amplify the sample. ND = none detected.

**Supplemental Table 2:**
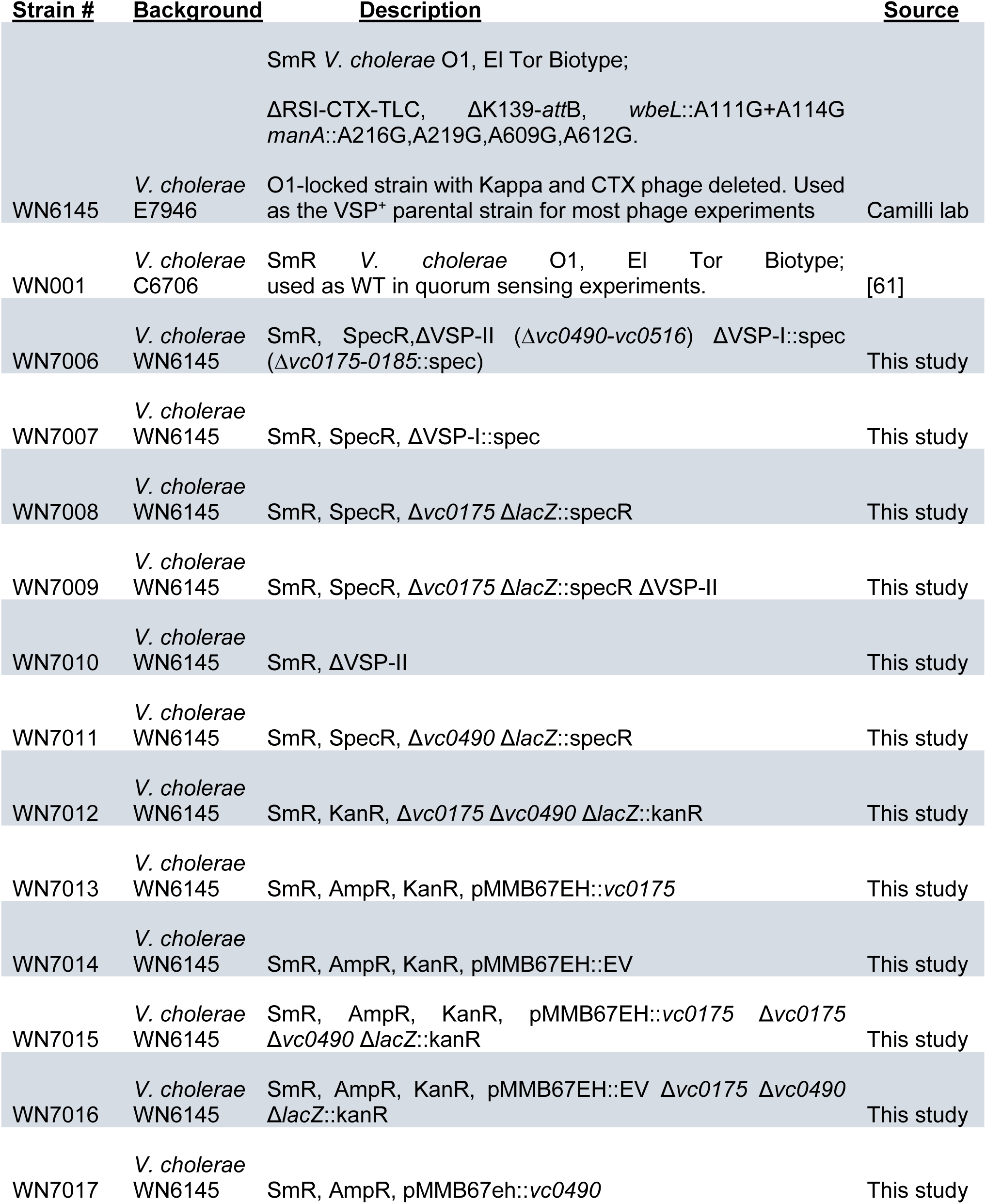
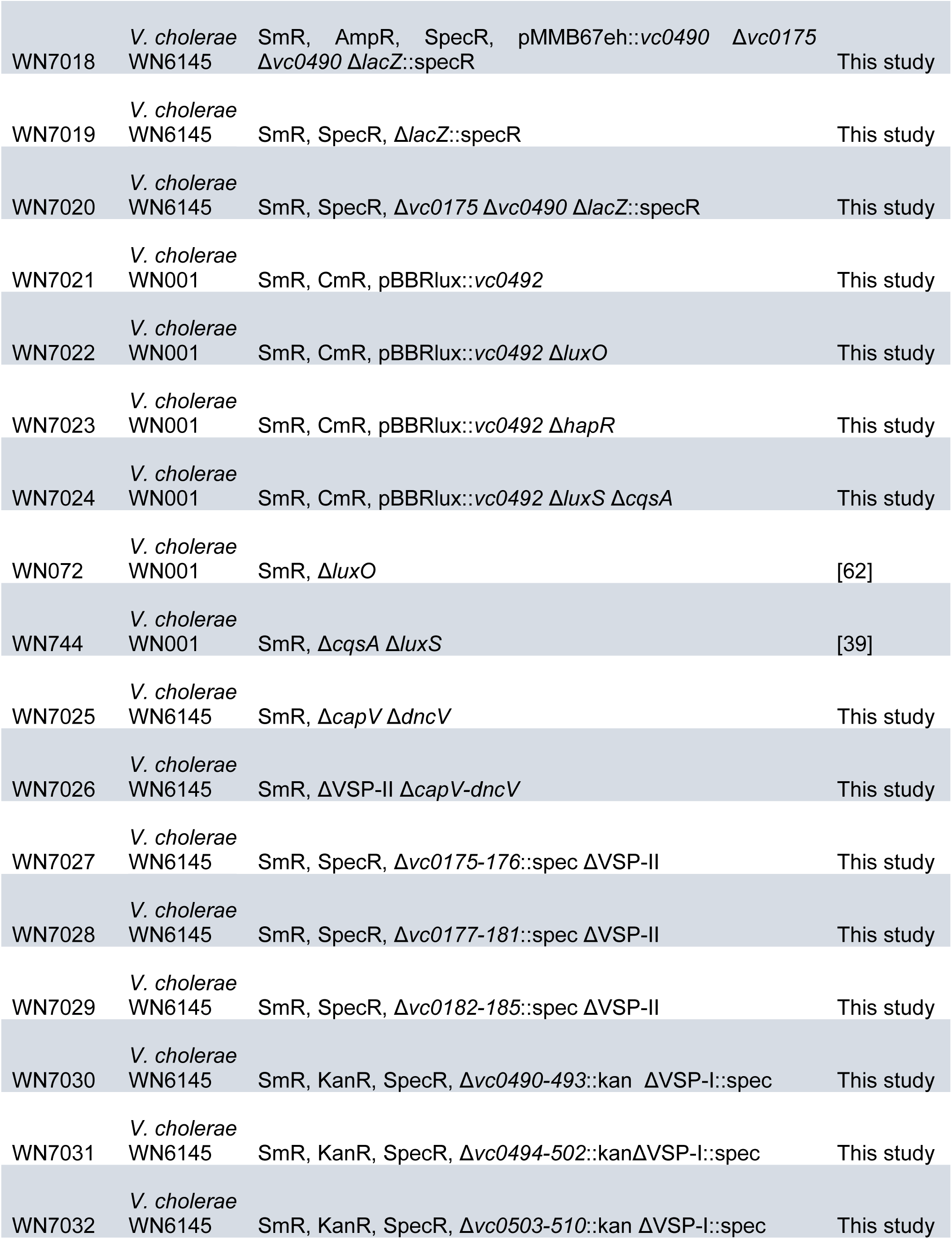
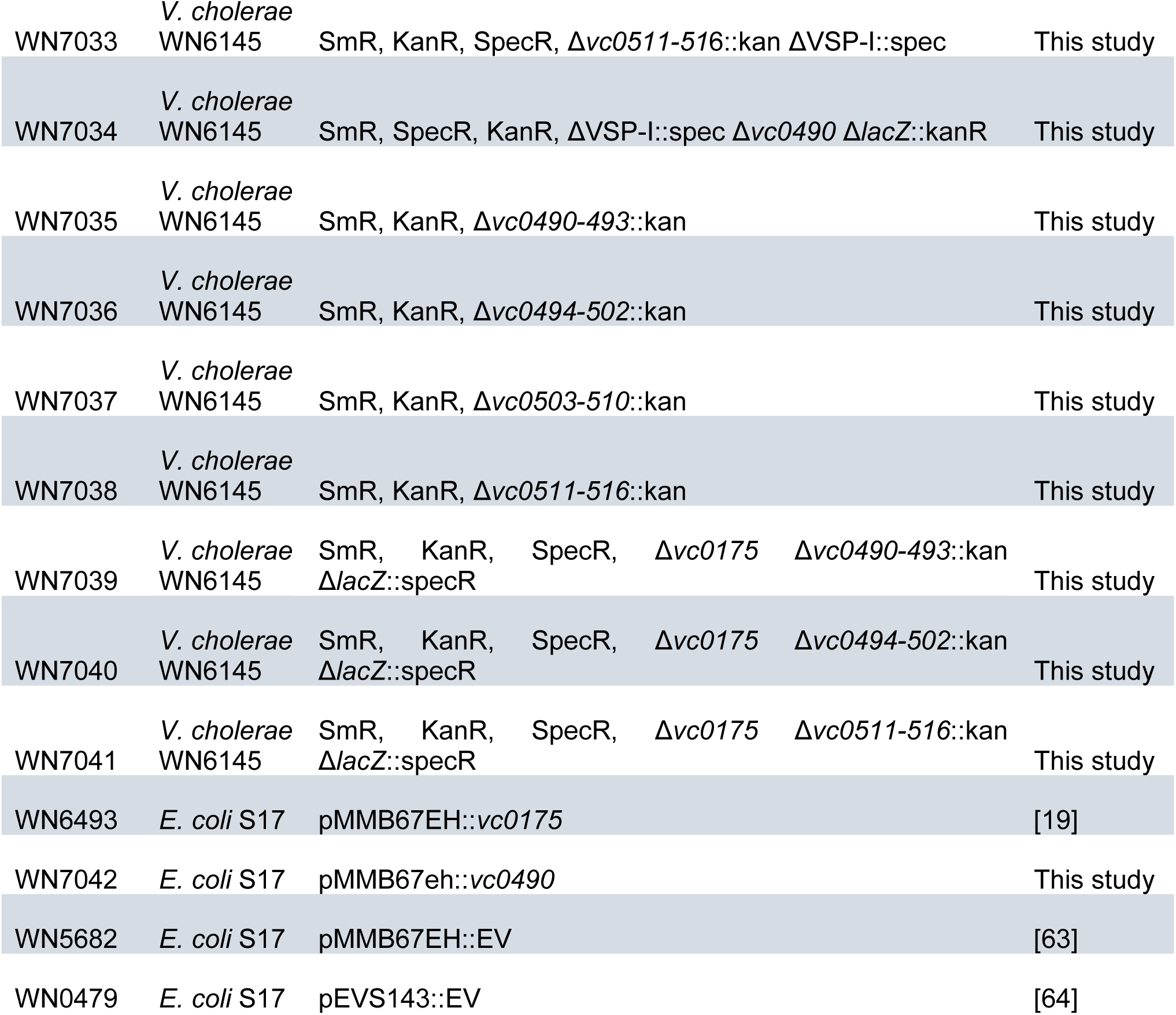
Bacterial strains used in this study

